# Working memory and decision making in a fronto-parietal circuit model

**DOI:** 10.1101/104802

**Authors:** John D. Murray, Jorge Jaramillo, Xiao-Jing Wang

**Affiliations:** Department of Psychiatry, Yale University School of Medicine, New Haven, CT 06510, USA; Center for Neural Science, New York University, New York, NY 10003, USA; NYU-ECNU Joint Institute of Brain and Cognitive Science, NYU Shanghai, Shanghai, China

## Abstract

Working memory (WM) and decision making (DM) are fundamental cognitive functions involving a distributed interacting network of brain areas, with the posterior parietal and prefrontal cortices (PPC and PFC) at the core. However, the shared and distinct roles of these areas and the nature of their coordination in cognitive function remain poorly understood. Biophysically-based computational models of cortical circuits have provided insights into the mechanisms supporting these functions, yet they have primarily focused on the local microcircuit level, raising questions about the principles for distributed cognitive computation in multi-regional networks. To examine these issues, we developed a distributed circuit model of two reciprocally interacting modules representing PPC and PFC circuits. The circuit architecture includes hierarchical differences in local recurrent structure and implements reciprocal long-range projections. This parsimonious model captures a range of behavioral and neuronal features of fronto-parietal circuits across multiple WM and DM paradigms. In the context of WM, both areas exhibit persistent activity, but in response to intervening distractors, PPC transiently encodes distractors, while PFC filters distractors and supports WM robustness. With regards to DM, the PPC module generates graded representations of accumulated evidence supporting target selection, while the PFC module generates more categorical responses related to action or choice. These findings suggest computational principles for distributed, hierarchical processing in cortex during cognitive function, and provide a framework for extension to multi-regional models.

## Introduction

Cognitive functions engage distributed networks of areas in the primate brain, with prefrontal cortex (PFC) and posterior parietal cortex (PPC) as key nodes (Duncan, 2010; Mitchell et al., 2016; Domenech et al., 2017). Working memory (WM) and decision making (DM) are fundamental building blocks of cognition that recruit a common prefrontal-parietal network, with WM- and DM- signals partially overlapping at the neuronal level (Meister et al., 2013). Both PPC and PFC exhibit characteristic neural activity of WM and DM. WM is associated with stimulus-selective persistent activity that spans the mnemonic delay (Goldman-Rakic, 1995; Constantinidis and Procyk, 2004). DM is associated with ramping dynamics reflecting the accumulation of evidence and target selection (Schall, 2001; Gold and Shadlen, 2007). The general similarity of neural activity of PPC and PFC during WM and DM has supported the view that they make comparable contributions to these functions.

Important open questions are to identify how PPC and PFC interact during WM and DM and what their specialized roles may be. For instance, WM-related persistent activity in these areas may be a locally generated phenomenon or, alternatively, the result of distributed inter-areal interactions (Christophel et al., 2017). Despite general similarities between neural responses in PPC and PFC, important differences, including in distractor processing and evidence accumulation, have been found that provide insight into their unique contributions to WM and DM (Katsuki and Constantinidis, 2012b; Suzuki and Gottlieb, 2013; Hanks et al., 2015). It is unclear to what degree function specialization of areas may be due to intrinsic differences in local microcircuitry (Murray et al., 2014b; Katsuki et al., 2014). In addition to differences between areas, single-neuron recordings have revealed a diversity of functionally-defined cell types within fronto-parietal circuits, across both WM and DM (Schall and Thompson, 1999; Ferraina et al., 2002; Lawrence et al., 2005). These findings raise questions of the division of labor among brain areas, or functional cell types, within distributed cortical circuits during cognition.

Biophysically-based computational models of cortical circuits have characterized neural circuit mechanisms for WM and DM functions. A class of models called *attractor networks* can perform these functions through strong recurrent synaptic interactions (Wang, 2008). In the attractor network framework, strong synaptic connections among neurons can provide reverberatory excitation, potentially mediated by slow NMDA receptors (Wang, 1999), that maintains a stimulus-selective persistent activity pattern for WM (Amit and Brunel, 1997; Wang, 2001; Machens et al., 2005). Strong lateral inhibition, mediated by GABAergic interneurons, can enforce selectivity of the WM representation, preventing an unstructured spread of excitatory activity (Compte et al., 2000; Brunel and Wang, 2001). Attractor networks can also perform slow integration and categorical, winner-take-all competition for perceptual DM (Wang, 2002; Wong and Wang, 2006). Indeed, strong recurrent excitation and lateral inhibition are required for winner-take-all DM in these models. Attractor networks therefore constitute a flexible type of ‘cognitive circuit’ capable of performing both WM and DM (Wang, 2013). In contrast to these theoretical advances in characterizing how local microcircuits can support cognitive processing, the computational principles for distributed cognitive processing in multi-regional cortical networks remain poorly understood.

To address these issues, we developed a biophysically-based computational model of two reciprocally connected modules, potentially representing circuits in PPC and PFC. We found that a single local circuit faces a tradeoff between optimization for WM vs. DM function. This performance tradeoff can be ameliorated in the distributed circuit, whose network properties are functionally desirable for both WM and DM. With a single set of network parameters, the distributed model can capture salient empirical observations from single-neuron recordings in PPC and PFC during WM and DM. In the context of WM, the model captures the relative roles of the two areas in distractor filtering. With regards to DM, the model captures key properties of functional cell types and the timing of their activity during visual search tasks. We propose that this cortical circuit model can provide insight into canonical features of distributed cognitive processing.

## Materials and Methods

### Model architecture

We constructed a distributed circuit model that is able to perform WM and DM computations. The model is comprised of two reciprocally interacting modules (Figs. 1 and 3A). Each module contains two selective, excitatory populations, labeled *A* and *B* (Wong and Wang, 2006; Wong et al., 2007). Within a module, the two populations have self-excitation and interact through a local inhibitory population that allows for cross-inhibition between the two excitatory populations. Each recurrently-connected excitatory population receives inhibition from a common pool of interneurons. Inhibition is linearized so that projections between the two excitatory populations *A* and *B* are effectively represented by negative weights (Wong and Wang, 2006). The two modules interact through long-range projections that are structured according to the stimulus selectivity of populations within each module. Long-range projections between modules are structured so that populations with the same selectivity are connected through excitatory projections whereas populations with different selectivity are connected via net inhibitory projections. The two modules are labeled 1 and 2 and external input related to the stimulus enters into Module 1 (Fig. 3A).

**Figure 1:**
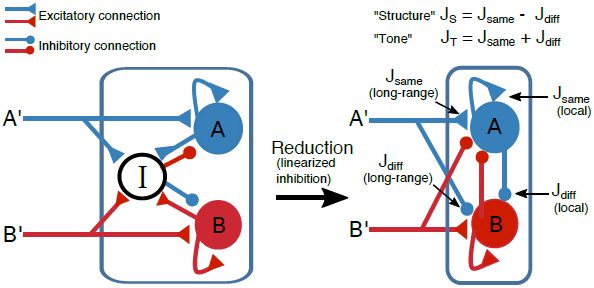
Circuit schematic of the firing-rate model. A module is defined as a set of two excitatory populations where each population is selective to one of two spatial, directional, or object stimuli (left). Each excitatory population is recurrently connected and also receives inhibition from a common pool of interneurons. The effects of inhibition and recurrent excitation are to generate bistability for WM and winner-take-all dynamics and ramping activity through slow reverberation for DM. Population A (B) receives input either from spatially-selective stimulus A’ (B’) or from another population A (B) in another module. The circuit dynamics can be simplified (right) by linearizing inhibition, so that effectively inhibition is represented by negative weights. Thus, the effect of the pool of interneurons is implicit in the inhibitory connections between the excitatory populations. In general, synaptic weights *J* can connect two selective populations of either the same (*J*_same_ > 0) or opposite (*J*_diff_ < 0) stimulus-selectivity and can be either local or long-range. The structure *J*_S_ = *J*_same_ — *J*_diff_ denotes the total recurrent strength while the tone *J*_T_ = *J*_same_ + *J*_diff_ denotes the net input onto a population. Synapses labeled with triangles and circles denote net-excitatory and net-inhibitory connections, respectively.

### Dynamics and stimuli

We constructed a population firing-rate model for each population *i* = *A, B* in the two modules. The firing-rate dynamics of the population *i* in Module *n* = 1, 2 are dominated by the dynamics of the average NMDA synaptic gating variable *S*^*n*^_*i*_. This approximation is based on the fact that the dynamics of the NMDA synaptic gating variable is slower than other time scales in the system (Wang, 2002; Wong and Wang, 2006; Wong et al., 2007). The gating variable *S*^*n*^_*i*_ is described by:

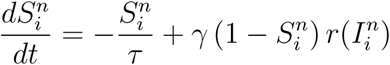
where the NMDA time constant τ = 60 ms, γ = 0.641 controls the rate of saturation of *S*, and *r*(*I*_*i*_^*n*^) is the firing rate of the population *i* as a function of the input current *I*_*i*_^*n*^. The firing rate as a function of input current is given by the F-I curve relation (Abbott and Chance, 2005):

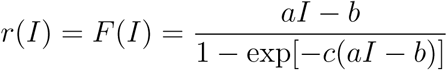
with *a* = 270 
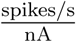
, *b* = 108 spikes/s, and *c* = 0.154 s. The input current to population *i* = *A, B* in Module *n* = 1, 2 is given by:

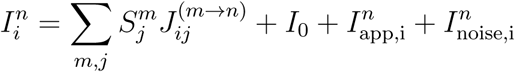
where **J*_ij_^(m→n)^* is the connection weight from population *j* in Module *m* to population *i* in Module *n*, *I*_0_ = 0.3347 nA is the background current, *I*^*n*^_app,i_ is the applied current to population *i* in Module *n* from external sources, and *I*^*n*^_noise,i_ is the noise current to population *i* in Module *n*.

In this study, the external current *I*_app_ is specified for three different task contexts: (1) WM with distractors, (2) perceptual DM with stochastic discrete input, and (3) perceptual DM or target selection in visual search. These currents will be specified below when we discuss the contexts and tasks.

The noise current to each population follows Ornstein-Uhlenbeck dynamics with the time constant of AMPA synapses:

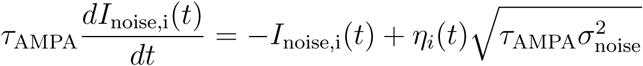
where τ_AMPA_ = 2 ms, η is Gaussian white noise with zero mean and unit variance, and σ_noise_ sets the strength of noise. As in Wong et al. (2007), we set σ_noise_ = 0.009 nA.

### Connectivity

The connectivity in our model is specified by the sign and magnitude of the connection weights between the selective excitatory populations, and whether the connections are local (within a module) or long-range (across modules). To this end, it is useful to express the connection weights with the terms:

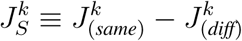

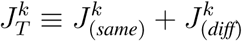
where *J*_(*same*)_ denotes the positive connection weight between same-selectivity populations, e.g. from population A in Module 1 to population A in Module 1 or 2. *J*_(*diff*)_ denotes the negative connection weight between different-selectivity populations, e.g. from population A in Module 1 to population B in Module 1 or 2, and *k* = 1 → 1, 1 → 2, 2 → 1, 2 → 2 defines whether the connection is local or long range. We define *J*_*S*_ as the *structure* of the network, since it reflects the magnitude of same-selectivity excitation and different-selectivity cross-inhibition and thus the total recurrent strength. Analogously, we define *J*_*T*_ as the tone of the network, which reflects the net input onto a particular population. Local connectivity for Module 1 is set by: *J*_*S*_^(1→1)^ = 0.35 nA and *J*_*T*_^(1→1)^ = 0.28387 nA. Local connectivity for Module 2 is set by: *J*_*S*_^(2→2)^ = 0.4182 nA and *J*_*T*_^(2→2)^ = 0.28387 nA. The only difference in local network properties between modules is that Module 2 has an enhanced structure as compared to Module 1: *J*_*S*_^(2→2)^ > *J*_*S*_^(1→1)^.

The decomposition of connectivity in terms of structure *J*_*S*_ and tone *J*_*T*_ is useful for capturing the impact of changes in activity that are symmetric or asymmetric between the populations of a module. The input current *I* to a given population from Module *m* to Module *n* can be written as (see first term in right-hand side of Eq.3)

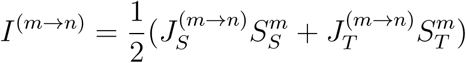
where *S* is similarly redefined through *S*_*S*_^*m*^ ≡ *S*(^*m*^_*same*_) — *S*^*m*^_(*diff*)_ and *S*_*T*_^*m*^ ≡ *S*(^*m*^_*same*_) + *S*^*m*^_(*diff*)_. When activity in a module is equal in the two populations (i.e., *S*_*S*_^*m*^ = 0), the net input from that module is determined only by the tone *J*_*T*_ and not by structure *J*_*S*_.

For both long-range projections between modules, we constrain them to have pathway-specific excitation/inhibition (E/I) balance:

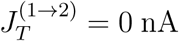

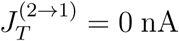

so that the excitatory weight to a given population is counteracted by inhibition of equal magnitude but opposite sign. If a pathway from one module to another exhibits balance (*J*_*T*_ = 0), the impact of one module to another is only nonzero when the populations have unequal activity, i.e., *S*_*S*_^*m*^ ≠ 0 (see also Vogels and Abbott (2009)). The structure of the projection from Module 1 to Module 2 is set by: *J*_*S*_^(1→2)^ = 0:15 nA. The structure of the projection from Module 2 to Module 1 is set by: *J*_*S*_^(2→1)^ = 0:04 nA. We can easily translate the structure *J*_*S*_ and tone *J*_*T*_ into individual synaptic weights. For example, *J*_*AB*_^(1→2)^ denotes the feedforward projection between the population *A* in the first module onto the population *B* in the second module and is given by:

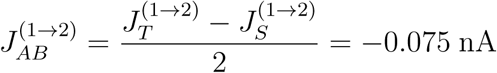

For the results in this study, we associate Module 1 with the PPC and Module 2 with the PFC.

### Working memory and decision-making in a local network

To characterize WM, we studied the generation of stimulus-selective persistent activity states and their robustness against intervening distractor inputs. In Fig. 2*C*,*D*, we characterized the robustness of WM against distractors in a local one-module circuit, over a two-dimensional parameter space of structure *J*_*S*_ and applied current *I*_*app*_. For these results, bifurcations and continuations were calculated using PyDSTool, a Python-based platform developed for the analysis of dynamical systems (Clewley, 2012). In Fig. 2*H* we defined the discrimination threshold *x*_disc_ as the minimum contrast to achieve 81.6% accuracy in a simulated two-alternative forced choice task. This number follows from the equation commonly used to fit psychometric curves

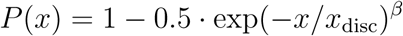
where *P* is the accuracy in the task, *x* is the contrast, *x*_disc_ is the discrimination threshold, and *β* determines the slope of the psychometric curve. We varied the recurrent structure *J*_*S*_ from *J*_*S*_ = 0.35 to *J*_*S*_ = 0.42 nA in steps of 0.01 nA and used Eq. 10 to fit the behavior of our simulated model and determine the discrimination threshold.

**Figure 2:**
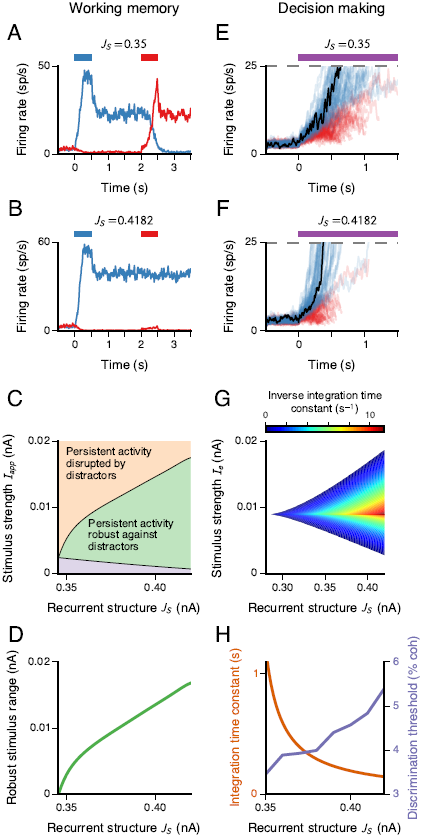
Tradeoffs between WM and DM function in a local attractor network model. A, Neural activity for a single WM trial. The colored bars mark presentation of input current to the population A (blue) and B (red), with strength *I*_app_ = 0.0295 nA. For a recurrent structure *J*_*S*_ = 0:35 nA, the circuit generates a stimulus-selective persistent memory state, but it is vulnerable to intervening distractors. B, At increased recurrent structure *J*_*S*_, WM activity in the circuit is robust against distractors. C, Robustness of WM as a function of recurrent structure *J*_*S*_ and stimulus strength *I*_app_. In the purple lower region, the stimulus is too weak for the target to induce a transition from the baseline state to the memory state. In the green middle region, the network can perform WM that is robust against intervening distractors. In the orange upper region, the stimulus current is strong enough for a distractor to disrupt target-related memory. D, WM robustness increases with increasing recurrent structure *J*_*S*_. The robust stimulus range is defined as the range of stimulus strength *I*_app_ in which persistent activity is robust against distractors, i.e. by the height of the green region in C. E, Neural activity during DM for zero-contrast stimulus (i.e., equal-strength input to both populations), for 100 trials in which the blue population first reached threshold. The colored bar marks stimulus presentation, with strength *I*_*e*_ = 0.0118 nA. The black trace marks the firing rate of the winning population for the trial with median reaction time. F, At increased recurrent structure *J*_*S*_, integration is shorter, limiting evidence accumulation, and ramping to threshold occurs sooner. G, Integration time constant as a function of recurrent structure *J*_*S*_ and stimulus strength *I*_*e*_ for a zero-contrast signal. The integration time constant is defined as the absolute value of the inverse eigenvalue of the unstable mode of the saddle point in the system (Wong and Wang, 2006). The inverse of the integration time constant is plotted. The white region marks where the symmetric state is stable, and therefore the network is not in a winner-take-all regime. H, DM performance degrades with increasing recurrent structure. For a fixed stimulus strength (here with *I*_*e*_ = 0.0118 nA), the integration time constant decreases with *J*_*S*_. Correspondingly, the discrimination threshold increases, indicating degraded performance.

**Figure 3:**
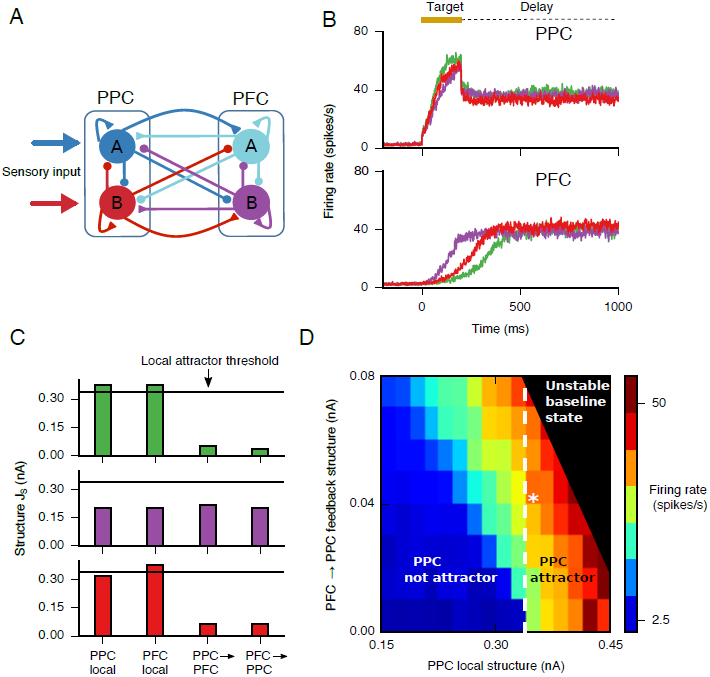
A distributed cortical model reproduces spatially selective persistent activity through local and long-range connections. A, The circuit is composed of two reciprocally connected modules, PPC and PFC, and each module consists of two excitatory neural populations selective to a stimulus A and B, respectively. The circuit model is endowed with self-excitation and cross-inhibition. Neurons in the PPC receive the sensory stimulus and convey the information to the PFC via long-range net-excitatory and net-inhibitory projections, as in Fig. 1. B-D, Local and long-range structure jointly contribute to persistent activity. B, PPC and PFC populations reach the same level of persistent activity in the steady state across the three scenarios depicted in C, demonstrating the joint contributions of long-range and local connectivity. Gold bar denotes stimulus presentation. C, Structure values reflecting local (within-module) and long-range (across-module) connectivity for three scenarios are shown: 1) PPC and PFC both independently support persistent activity (green, top), 2) neither PPC nor PFC is capable of persistent activity independently (purple, middle), and 3) only PFC independently supports persistent activity (bottom, red). Black horizontal line denotes the threshold for a local module to support persistent activity independently (i.e., multiple stimulus-selective attractor states). D, Steady-state firing rate for the activated population of the PPC module in the memory state, as a function of PPC local structure and PFC→PPC feedback. The PFC local and feedforward PPC→PFC structure are fixed. In the region in the upper-right corner, the baseline state is unstable. In the region to the right of the white dashed line, the PPC is an independent attractor. The white asterisk marks the parameter values used for the WM and DM simulations in Figs. 4–8.

In Fig. 3, to understand how the dynamics of the distributed two-module circuit depends on the structure, we parametrically varied the Module 1 (PPC) local structure *J*_*S*_^(1→1)^ from 0.15 to 0.45 nA and the PFC→PPC feedback structure *J*_*S*_^(2→1)^ from 0 to 0.08 nA, in steps of 0.02 nA, while keeping all other parameters constant.

### Working memory and distractors in the PPC-PFC circuit

For Figs. 4 and 5, we simulated a WM task with distractors, based on a primate electrophysiology study using an visuospatial WM task in which a subject must hold in WM the position of a target, and filter intervening distractor stimuli appearing at other positions during the delay period (Suzuki and Gottlieb, 2013). We implemented a discrete version of this task with selectivity to two stimuli. A flash of 100 ms appears on one of two positions of a screen indicating the target position. The target to be held in WM, is the first stimulus presented. We set the target as a current *I*_app,A_ = *I*_target_ of 100-ms duration that is applied to population *A* in the PPC module. Distractors are defined as inputs *I*_app,B_ = *I*_distractor_ of equal duration applied to population *B* arriving after the target and at an opposite location of the visual field. Similar to (Suzuki and Gottlieb, 2013), we considered four times of distractor onset asynchrony (TDOA) defined as the onset of the distractor relative to onset of the target: 100, 150, 200, and 300 ms. Targets and distractor amplitudes were sampled randomly and independently from a Gaussian distribution with mean 0.09 nA and standard deviation 0.04 nA.

**Figure 4:**
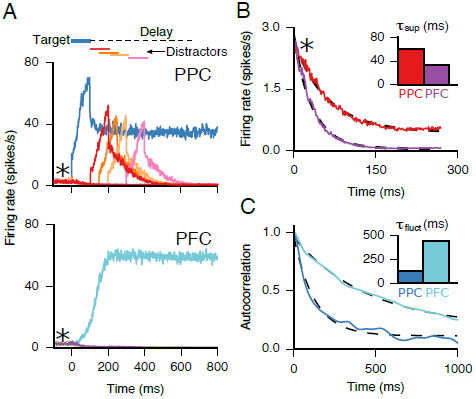
Firing-rate dynamics in the PPC-PFC circuit during a WM task. *A*, Top, the blue trace shows the response of the target-selective neural population in the PPC in response to a target presented at *t* = 0 ms, with no distractors. Red, orange, light orange, and pink traces show the response of the distractor-selective population to distractors presented at *t* = 100, 150, 200, 300 ms, respectively. Bottom, the cyan trace shows activity of the target-selective population in the PFC that receives the stimulus indirectly through the long-range projections from the PPC, in the no-distractor condition. The other traces show the distractor-selective population in response to distractors, as for PPC. However, these responses are not visible due to the strong filtering by surround suppression within PFC. *B*, Temporal dynamics of the two suppressed populations in PPC and PFC around the time of stimulus presentation (asterisks in *A*). *C*, Autocorrelation of the spontaneous firing rate shows the difference in fluctuation time scales τ_fluct_ between the PPC and PFC. Firing rate traces were smoothed with Gaussian window of width 20 ms before calculating the autocorrelation.

In Fig. 4*B*, we calculated the differences between the two modules in terms of how distractors are suppressed. For this we fitted the time courses of the firing rates *r*(*t*) of the populations selective to the distractor for each of the modules with an exponential function:

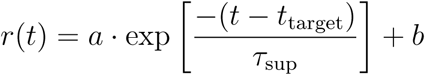
where *t*_target_ marks the time of target onset and also suppression of the distractor, τ_sup_ is the time scale of suppression, and *a* and *b* are parameters of the fit.

In Fig. 4 *C*, we performed an autocorrelation on the firing rate to reveal the intrinsic or fluctuation time scales of spontaneous activity. The firing rate was first filtered with a Gaussian function with window σ_filter_ = 20 ms. To compute the autocorrelation of the firing rate, we substracted the mean from the firing rate and then normalized. We then used Equation 11 to fit the normalized firing-rate autocorrelation and extract the respective time scales τ_fluct_.

We plotted error rates vs. time of distractor presentation relative to the target in Fig. 5 *C*, where an error was recorded when the population selective to the distractor is at the high memory state at *t* > 3000 ms.

**Figure 5:**
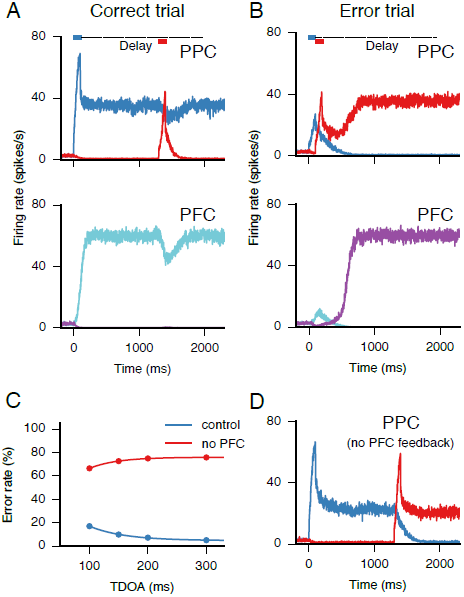
Relationship between neural dynamics and behavioral performance during a WM task with distractors (colors correspond to the schematic in Fig. 3A). *A*, Example of a correct, not distracted, trial. The target-selective population in PPC encodes the target in WM following stimulus onset at *t* = 0 ms (top, blue), while the distractor-selective PPC population transiently but strongly encodes the distractor following its presentation at *t* = 1300 ms (top, red). After distractor offset, feedback from the PFC switches the PPC back to encoding the target, enabling a correct response at the end of the trial. The PFC (bottom) is activated by the PPC’s response to the target, which is maintained in WM by the target-selective population in the PFC as well. Distractor presentation causes a transient suppression of the delay activity in the neurons encoding the target (cyan), but the distractor is not represented strongly (magenta) as it is in the PPC. *B*, Example of an error, distracted, trial. If the target precedes the distractor by a short interval (100 ms in this example), there is an increased probability of the distractor representation overriding the target representation, so that the distractor is encoded in persistent activity in both PPC and PFC (top, red; and bottom, purple). *C*, Simulated behavioral performance as a function of time of distractor onset relative to target (target-distractor onset asynchrony, TDOA). Distractibility decreases with longer TDOA (blue). Simulated lesion of PFC greatly increases distractibility (red). *D*, Effects of removal of PFC!PPC feedback. Absence of PFC feedback onto the PPC forces the PPC to encode the last presented stimulus, leaving it vulnerable to distractors.

### Evidence accumulation in the PPC-PFC circuit

As shown in Fig. 6, we simulated a simple version of a discrete evidence accumulation task based on a two-alternative forced choice task with perceptual decisions that rely on evidence accumulation from discrete auditory stimuli (Brunton et al., 2013). The auditory input consists of a sequence of clicks on both sides (left and right), parametrized by click frequency in units of clicks per second. For example, 10:24 constitutes a trial where 10 represents the click frequency for the left side, and 24 constitutes the click frequency for the right side. In the task the subject is rewarded for reporting which side (left vs. right) had the higher frequency signal. This task can be solved by integrating evidence, where each click represents a unit (quantum) of evidence (Hanks et al., 2015).

**Figure 6:**
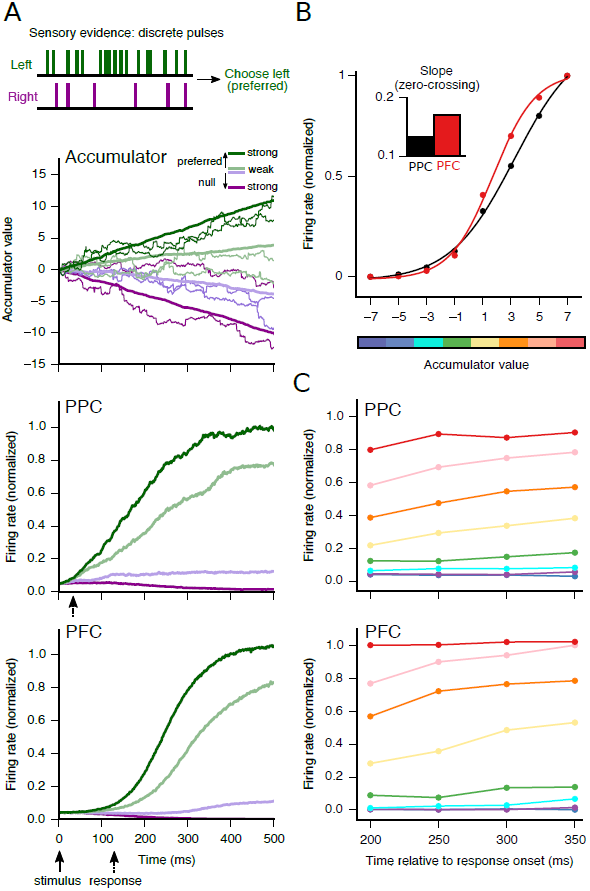
PPC and PFC differentially encode accumulated evidence during perceptual DM. *A*, The theoretical accumulator (top), PPC (middle), and PFC (bottom) integrate sensory evidence as a function of time and trial difficulty. Thick traces show the average over 60 trials for each difficulty condition while thin traces in the accumulator show single trials. *B*, The firing rate vs. accumulator relationship is more categorical, with a steeper slope at zero accumulator value, in the PFC than in the PPC, which has a more graded coding. The slopes at zero crossing are 0.13 and 0.17 for PPC and PFC, respectively (see also Hanks et al. (2015)). *C*, The relationship between firing rate in PPC and PFC and accumulator value as a function of time is stable. The eight accumulator values (from purple to red) correspond to the horizontal axis in *B*.

In our model, clicks are represented by a set of Poisson-distributed times, parametrized by rate and side of origin, either left or right. The rates for each side are such that they add to 34 clicks/s in total. For example, a difficult trial is 18 clicks/s:16 clicks/s while an easy trial is 30:4. The click times *t_i_* for each side are convolved with a current pulse kernel II of amplitude 0.0118 nA and pulse duration 50 ms, so that the current *I*_L,R_ is:

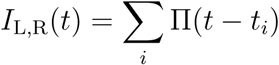

These currents, corresponding to left and right clicks, are fed onto the corresponding selective populations (left or right) of PPC, as well as to the accumulator which we now define. The accumulator is an implementation of a drift-diffusion model (Ratcliff, 1978), with parameters for drift, noise, and input stimulus (Brunton et al., 2013). Time evolution of the accumulator value *a* at time *t* is given by:

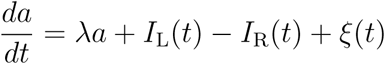
where λ is the drift and ξ(*t*) is Gaussian white noise with zero mean and standard deviation 0.1. We set λ = 0, corresponding to leak-free integration, for the simulations in Fig. 6, but small values of |λ| did not significantly alter these plots.

To obtain the relationship between an accumulator and PPC/PFC firing rates, we selected four time points relative to response onset in the PPC/PFC (*t* = 200, 250, 300, 350 ms) and obtained the distribution of firing rates and accumulator values for each of those time points (Hanks et al., 2015). For each of the time points, we binned the accumulator values from −7 to 7, with bin size of 2 and calculated the mean firing rate for each bin. Applying this procedure to the four selected time points, we obtain Fig. 6*C*, where normalized firing rate is shown as a function of time, and color coded as a function of accumulator value. Finally, to obtain the relationship between firing rate and accumulator value for the PPC and PFC, we averaged the firing rate over time for each of the accumulator values (Fig. 6*B*).

### Inputs and neural measures for a visual search task

In Figs. 7 and 8 we simulated a perceptual DM task analogous to target selection in visual search (Sato et al., 2001). Each module contains two populations that are selective to a target and a distractor, respectively. As in the WM paradigm, external stimuli enter as currents into the PPC module. These applied currents reflect the external stimulus as:

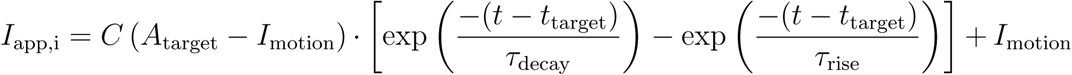

where 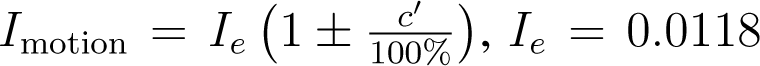, *I*_*e*_ = 0.0118 nA scales the overall strength of the input and *c*^′^, referred to as the contrast, sets the bias of the input for one population over the other (equivalent to the coherence in Wong and Wang (2006)), *A*_target_ and *t*_target_ determine the amplitude and the onset of the target, respectively, the time constants τ_decay_ and τ_rise_ determine approximately the decay and rise of the target-induced transient response and *C* is a normalization factor. A zero-contrast stimulus applies equal input *I*_*e*_ to each population in Module 1 (PPC, also denoted as Selection cells). In all of the simulations and when *c*^′^ > 0, the target-selective population receives the greater biased input. Due to noise, however, this does not guarantee that the target population will win, especially for low-contrast values.

**Figure 7:**
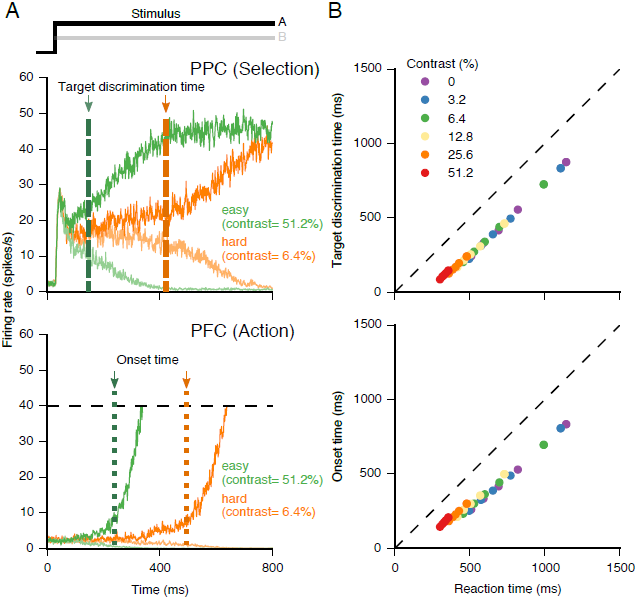
Dynamics of the distributed circuit model during a visuo-spatial DM task. *A*, (Top) Target and distractor cells in the Selection module receive stimulus inputs, integrate perceptual evidence, and discriminate the inputs (marked by target discrimination time). (Bottom) Following target discrimination in the Selection module, the corresponding population of Action cells is activated and begins ramping (marked by onset time). When one of the Action populations reaches a threshold of 40 spikes/s (black dashed lines), an overt response is triggered and a reaction time is registered. Green (orange) traces correspond to easy (hard) trials. Target and distractor cells are shown in thick and thin lines, respectively. Dashed lines mark the target-discrimination time in Selection cells defined as the time when the difference in firing rate of the two populations has reached 12 spikes/s. Dotted lines mark the onset time in Action cells defined as the time when the firing rate of one of the populations has reached 7 spikes/s. *B*, Target discrimination times in the Selection module (top) and onset times in the Action module (bottom) correlate with reaction times, both across and within contrast conditions. Reaction times were split into quintiles for each contrast level. Only correct trials are shown.

Because our model provides instantaneous firing rates for a population, we can define measures of neural activity directly for individual trials. For DM simulations (Figs. 7 and 8), we define and calculate the reaction time as the time at which the firing rate of a population in Module 2 (PFC, also denoted as Action cells) crosses a threshold (Hanes and Schall, 1996). We measure the discrimination time for Selection cells in Module 1 (PPC) through a threshold on the absolute difference between firing rates between the two populations. We measure the onset time for Action cells in Module 2 through a threshold on firing rate for the winning population. The thresholds for reaction time, discrimination time, and onset time at 40, 12, and 7 spikes/s, respectively. Reaction times are divided into quintiles for each of the contrast conditions. This is analogous to the division into short, intermediate, and long reaction time groups (Sato et al., 2001).

**Figure 8:**
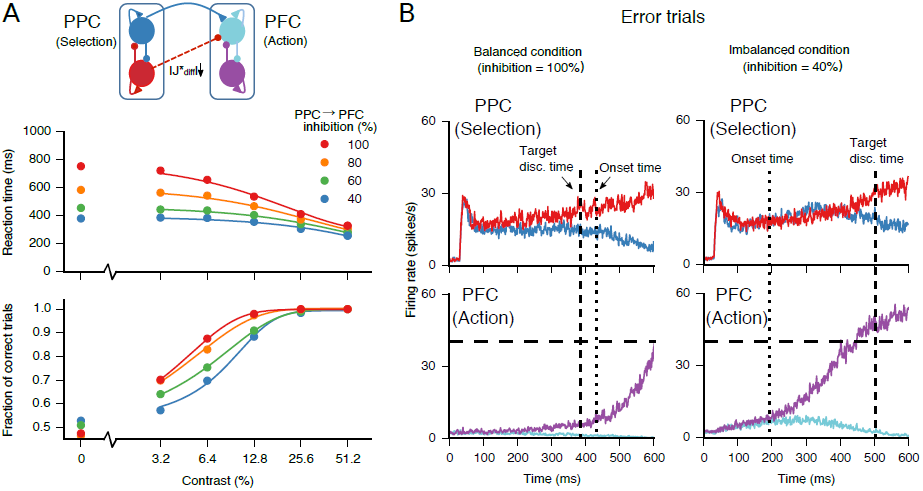
Pathway-specific excitation-inhibition balance disruption and speed-accuracy tradeoff. The degree of balance disruption is quantified by the percentage of the total inhibition required to balance excitation to a cell (100% corresponds to full balance). *A*, Speed-accuracy tradeoff. Reaction times decrease both as a function of contrast and degree of balance disruption (top). The fraction of correct trials increases as a function of contrast but decreases as a function of the degree of balance disruption (bottom). *B*, Error trials with (left) and without (right) pathway-specific balance. For balanced trials (100% inhibition), errors are due to mis-selection from cells in the Selection module and subsequent ramping of a population of cells in the Action module (top). For imbalanced trials, i.e., with excitatory bias (40% inhibition), errors can be due to early ramping in cells in the action module before any divergence has begun in the Selection module (bottom). Dashed black lines mark the target-discrimination time in the Selection module, and dotted black lines mark the onset time in the Action module.

### Disruption of pathway-specific excitation-inhibition balance

In Fig. 8 we examined the effects of pathway-specific excitation-inhibition balance in inter-Module projections. To this end, we systematically decreased the inhibitory weights of the projections from Module 1 to Module 2 as to alter the E/I ratio. The altered inhibitory weights *J*^*^_(diff)_ are

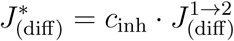
where *J*^1→2^_(diff)_ is the original, i.e., unaltered, inhibitory synaptic weight between population A (B) in Module 1 and population B (A) in Module 2 and *c*_inh_ ϵ {0.2,0.4,0.8,1}. According to Eq. 15, *c*_inh_ = 1 corresponds to balanced excitation and inhibition onto both populations in Module 2 while *c*_inh_ < 1 corresponds to imbalance, i.e., a net positive weight onto the populations (see Eqs. 6 and 8).

We fit the reaction time vs contrast relation in Fig. 8*A*, top, using an exponential function

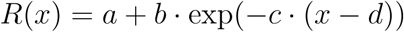
where *R* is reaction time, *x* is the contrast and *a, b, c* and *d* are free parameters of the fit. Similarly, we fit the fraction of correct trials vs contrast relation in Fig. 8*A*, bottom, with a sigmoid

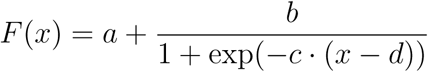
where *F* is the fraction of correct trials, *x* is the contrast and *a, b, c* and *d* are free parameters of the fit.

## Results

We have designed and characterized a distributed circuit model that supports persistent activity for WM and slow integration over time and winner-take-all competition for DM. The model is comprised of two reciprocally connected modules that model the posterior parietal cortex (PPC) and pre-frontal cortex (PFC). Each module consists of two populations of excitatory neurons and each population is selective to one of two spatial, directional, or object stimuli (see Fig. 1 and Materials and Methods). Local connectivity, i.e., connectivity within a module, is specified by recurrent excitation and cross-inhibition. Long-range connectivity, i.e., connectivity across modules, is specified by feedforward and feedback projections that are net-excitatory between same-selectivity populations and net-inhibitory between different-selectivity populations. Model parameters were chosen so that the same architecture and parameter set could capture several important neurophysiological dynamics observed in both WM and DM.

### Tradeoffs in working memory and decision making for a local circuit

We first characterized WM and DM function in a local, one-module network. Prior studies have shown that a local attractor network can perform both WM and DM (Wang, 2002; Wong and Wang, 2006). However, it has been less studied how well the same network can perform both functions, or what tradeoffs exist between optimization of local circuit properties for WM vs. DM (Standage and Paré, 2011). Of particular interest is the role of the local recurrent structure, parameterized by *J*_*S*_, here defined as the total recurrent strength including self-excitation within a population and cross-inhibition between populations (see Eq.5 in Materials and Methods).

#### Working memory and robustness against distractors

In the attractor network framework, the key requirement for WM function is multistability, i.e., the coexistence of multiple stable fixed points (attractor states) in the absence of a stimulus (Wang, 1999; Brunel and Wang, 2001). In the absence of a stimulus, the simplified two-population network studied here supports three stable states: one symmetric baseline state and two asymmetric memory states (Wong and Wang, 2006). Before stimulus onset, the network is in the symmetric baseline state with both populations, A and B, at low activity (Fig. 2*A*,*B*). After one population is sufficiently activated, it is able to remain persistently in a stimulus-selective, high-activity state in the absence of a stimulus.

In addition to maintenance over time, robust WM requires shielding internal representations from interference by both internal noise and external distraction. There is evidence that PPC and PFC have different susceptibilities for distractors to disrupt persistent activity. In general, PFC exhibits persistent activity that is robust against distractors, whereas posterior association areas in the temporal and parietal lobes exhibit persistent activity that is disrupted by distractors (Miller et al., 1993, 1996; Constantinidis and Wang, 2004; Qi et al., 2010; Suzuki and Gottlieb, 2013). Motivated by these findings, we explored the mechanisms of robustness against distraction and how they depend on network structure. In our model, distractors can be modeled as an intervening input during the WM delay to a different population than the one activated by the WM target. For instance, if population A is active in the memory state, a distractor is modeled as a subsequent input to population B.

We found that recurrent structure *J*_*S*_, i.e., the total self-excitation and cross-inhibition within the one-module network, plays a crucial role in determining the robustness against distractors in the local network (Fig. 2*A*–*D*). With moderate structure, the network generates WM-related persistent activity, but it is distractible. As shown in Fig. 2*A*, the distractor input switches the network to representing the distractor (red) instead of the target (blue). With high structure, in contrast, the distractor is filtered out and the target representation is robustly maintained (Fig. 2*B*). The network mechanism for robustness results from the combined effects of lateral inhibition and recurrent excitation set by *J*_*S*_ (Brunel and Wang, 2001). When population A is in the high-activity state, distraction requires both activation of population B and deactivation of population A. Activation of population B by the distractor input is counteracted by lateral inhibition. Recurrent excitation within population A counteracts the ability of inhibition from population B to deactivate it.

A dynamical systems analysis can formally characterize robustness of WM to distractors, through the bifurcation diagram defining how the system’s fixed points change as a function of input current to one population *I*_app_. The bifurcation diagram gives the range of input current strengths in which WM is robust against distractors. We can examine how this range varies as a function of recurrent structure *J*_*S*_. Fig. 2C,D shows how this robustness range of *I*_app_ changes with *J*_*S*_. We found that the distractibility threshold (boundary between green and orange regions) increases with *J*_*S*_. The memory induction threshold (boundary between purple and green regions) decreases as *J*_*S*_ increases, widening the robustness range further. The net effect is that the robust stimulus range increases with higher recurrent structure *J*_*S*_ (Fig. 2D). This analysis suggests that increased robustness against distractors in PFC, compared to PPC, may be due to higher network structure (Brunel and Wang, 2001).

#### Decision making and slow integration of evidence

We now consider the ability of the attractor network to perform perceptual DM functions. In the attractor network framework, two key requirements for DM are slow accumulation of evidence over time and winner-take-all competition (Wang, 2002; Wong and Wang, 2006). For a zero-contrast stimulus, the two populations receive equal input, differing only through the noise term. The network nevertheless performs categorical selection through one population going to a high-activity state and the other to a low-activity state.

To subserve perceptual DM, the network should be able to integrate evidence over time when the signal-to-noise ratio is low (Gold and Shadlen, 2007). In the model, a decision is made when the corresponding neural population reaches a threshold firing rate. As shown in Fig. 2*E*,*F*, the recurrent structure *J*_*S*_ plays a crucial role in determining the timescale of integration as reflected in ramping neural activity, here with a zero-contrast stimulus. With moderate structure, the network ramps relatively slowly (median decision time ~560 ms), indicating the network implements slow integration of sensory evidence. With higher recurrent structure, the network ramps substantially faster (median decision time ~ 430 ms), indicating a more limited duration over which the network integrates sensory evidence.

A dynamical systems analysis can formally characterize the integration timescale. In response to a zero-contrast stimulus, the network’s symmetric state is a saddle point in the (*S*_1_; *S*_2_) phase plane (Wong and Wang, 2006). The timescales associated with this saddle point, along with the strength of noise, largely determine the timescales of integration. The antisymmetric mode, whose positive eigenvalue indicates that it is the unstable mode, is the direction in which integration occurs and which leads to categorical choice. The integration timescale τ_int_ can be defined as the inverse of this eigenvalue. Fig. 2G shows the dependence of τ_int_ on network structure *J*_*S*_ and stimulus current strength *I*_*e*_, for zero-contrast stimulus. As the system approaches the bifurcations that form the boundaries of the winner-take-all regime, the integration timescale increases toward infinity. For a fixed *I*_*e*_ in a winner-take-all DM regime (Fig.2G), increasing *J*_*S*_ decreases the integration timescale, which limits the duration over which perceptual integration can occur. If less evidence is integrated, we expect to have more errors for a fixed contrast (coherence in Wong et al. (2007); Gold and Shadlen (2007)) value. This degradation in performance is measured by the increase of the discrimination threshold, defined as the contrast necessary to achieve a predefined performance level, here 81.6 % correct (see Eq. 10 in Materials and Methods). Indeed, the shortened integration timescale degrades DM performance, as reflected in the discrimination threshold from the psychometric function (see Fig. 2H). Therefore, although recurrent structure must be above a threshold value in order for the network to perform winner-take-all selection, further increasing recurrent structure limits the gradual accumulation of sensory evidence.

These findings illustrate a tension between WM and DM function in a local attractor network. There is a tradeoff between the two as recurrent structure **J*_S_* is varied: higher structure increases robustness against distractors for WM (Fig. 2*D*), but at the expense of shortening the integration timescale which degrades DM performance (Fig. 2*H*). As we show and discuss below, this performance tradeoff may be ameliorated in a distributed circuit in which local modules have different strengths of recurrent structure.

### Persistent activity in a circuit model with local and long-range connections

Visual WM recruits persistent activations that are distributed across PPC and PFC (Chafee and Goldman-Rakic, 1998), which are mediated by PPC-PFC interactions (Chafee and Goldman-Rakic, 2000; Ferraina et al., 2002; Salazar et al., 2012; Dotson et al., 2014). The distributed circuit model we developed is composed of two reciprocally connected modules that can support persistent activity independently (Fig. 2A,B). What are the roles of local (within-module) vs. long-range (across-module) connections in supporting persistent WM states in this distributed circuit model? We model visual stimuli as inputs to PPC (Module 1, Fig. 3*A*), following the dorsal visual pathway (Felleman and Van Essen, 1991) and the ordering of activations during bottom-up visual processing as well as during target selection (Buschman and Miller, 2007; Ibos et al., 2013; Siegel et al., 2015), although some experiments suggest that input can also rapidly reach PFC through other pathways (Katsuki and Constantinidis, 2012a). Fig. 3*B* shows the model circuit response to a stimulus input to one PPC population for different values of the recurrent structure *J*_*S*_ (Fig. 3*C*). PPC responds vigorously to the stimulus, and propagates this signal to PFC. Following the offset of the stimulus, i.e., during the WM delay, both PPC and PFC encode the stimulus through selective persistent activity. On the basis of WM delay activity alone, PPC and PFC therefore have similar WM activity, as observed experimentally (Chafee and Goldman-Rakic, 1998; Qi et al., 2010; Suzuki and Gottlieb, 2013).

We examined the roles of local and long-range structure in determining persistent activity. Figure 3*B*,*C* show how different combinations of local and long range parameters can give rise to similar delay activity in the PPC and PFC modules. A transient stimulus input can switch the state of the circuit from a low-activity baseline state to a persistent and selective high-activity memory state. The stable and persistent delay activity is a reflection of the attractor dynamics in the combined PPC-PFC circuit. Interestingly, these attractor dynamics are present in the combined circuit even if the individual modules are not attractor networks independently (Fig.3*C*). This demonstrates that in a distributed network, the observation of persistent activity in a local area does not necessarily imply that the area is independently capable of multistability. We then examined the relationship between the firing rate, memory states, and structure in the distributed circuit. In Fig. 3*D*, the tradeoff in local and long-range structure is made explicit for the PPC firing rate in a high memory state. In conclusion, persistent activity in the PPC-PFC circuit is supported by both local and long-range structure, with the persistent activity states determined by the total, i.e., combined local and long-range, structure.

### Working memory with distractors in the parietal-prefrontal circuit

The ability for a WM circuit to encode and maintain information robustly while filtering out distractors is crucial for WM function (Sakai et al., 2002; Suzuki and Gottlieb, 2013). Our distributed cortical model is able to selectively encode a target stimulus in WM in the presence of distractors (Figures 4 and 5). Within the PPC and PFC modules, one neural population is selective to the target, and the other to the distractor. Fig. 4A shows how the distributed circuit responds to target and distractor stimuli during WM. Presentation of the target stimulus activates the selective population in PPC, which transmits this information to the PFC module via the feedforward long-range projections. Following stimulus withdrawal, the target is encoded in PPC and PFC in persistent activity. An intervening distractor, presented during the delay, competes with the mnemonic target representation in the network activity (Fig. 4A). If WM is robust, the distractor’s response is transient and target-coding persistent activity is maintained.

Several notable observations of distractor processing in the model are in line with single-neuron recordings from PPC and PFC during visuospatial WM. First, distractor responses are weaker than the target response (Suzuki and Gottlieb, 2013; Falkner et al., 2010; Zhang et al., 2017). This surround suppression is mainly due to the cross-inhibition from the active target population to the distractor population. Second, the peak distractor amplitude also decreases as the time of distractor presentation relative to target increases (Suzuki and Gottlieb, 2013). This results from the network dynamics of the synaptic gating variables. At shorter distractor onset times, the synaptic gating variables of the target and distractor populations have not reached their steady-state levels, i.e., a high value for the target population and a low value for the distractor population. Therefore, the suppressive effect from the target population on the distractor responses will slightly increase over time. Finally, and most strikingly, distractor responses are markedly different in PFC as compared to PPC: PPC represents the distractor strongly during its presentation (Suzuki and Gottlieb, 2013; Powell and Goldberg, 2000; Falkner et al., 2010), while PFC strongly filters distractors (Suzuki and Gottlieb, 2013; Everling et al., 2002). This is because the PFC module has a higher local structure *J*_S_ and thus stronger recurrent dynamics than the PPC. The stronger recurrence makes target encoding in PFC more robust against transient distractor inputs, which are effectively filtered. From the dynamical systems perspective, the PFC module has a wider basin of attraction than the PPC (Fig. 2) (Brunel and Wang, 2001; Wong and Wang, 2006). To summarize, the transient encoding of distractors is weaker than target encoding, and weaker in PFC than in the PPC.

Single-neuron recordings have found that WM activity in PPC and PFC generates surround suppression even on baseline activity, in the absence of distractors (Suzuki and Gottlieb, 2013; Falkner et al., 2010; Funahashi et al., 1989). We analyzed differences between the PPC and PFC modules in terms of the dynamics of this surround suppression during target encoding (Fig. 4B). Relative to PPC, in PFC surround suppression of the distractor population is (1) stronger, i.e., towards a lower baseline activity, and (2) more rapid, i.e., with lower time constant. These features are non-trivial, given that the stimulus directly drives PPC, whereas PFC is activated via projections from PPC. They result from the higher structure in PFC relative to PPC. Both of these features are in line with single-neuron features of distractor suppression (Suzuki and Gottlieb, 2013).

The timescale of surround suppression can be contrasted to the intrinsic timescale of activity fluctuations in the baseline state. Primate cortex shows a hierarchical organization of this dynamical feature, with intrinsic timescales increasing along the cortical hierarchy, with dorsolateral PFC exhibiting a longer timescale than posterior parietal area LIP (lateral intraparietal area) (Murray et al., 2014a). We examined whether this hierarchical organization was consistent with the distributed circuit model we developed. To this end, we calculated the autocorrelation of spontaneous fluctuations in the PPC and PFC, and found that PFC has a longer time constant indicating a slower decay (Fig. 4C), in line with empirical findings (Murray et al., 2014a). Notably, this ordering of intrinsic timescale (longer in PFC than PPC) is opposite to the ordering of the surround-suppression timescale (shorter in PFC than PPC). These contrasting time-scale orderings suggest that the difference between PPC and PFC is not simply that one area is more sluggish than the other one, but rather that these dynamics arise from interactions in the distributed circuit.

### Distractor-induced errors in working memory

We now examine the circuit’s dynamics in relation to behavioral performance on a WM task with distractors (Fig. 5). An example of a correct trial is shown in Fig. 5A. The PPC first encodes the target presented at *t* = 0 ms which is subsequently encoded in persistent activity in both PPC and PFC, and after 1300 ms a distractor is presented to another population in the PPC. During distractor presentation, the distractor-selective population in PPC encodes the distractor strongly in its firing rate, while persistent activity in the target-selective population is transiently and mildly perturbed. PPC’s strong distractor response is due to two features of the model: stimulus input into PPC is strong and PPC is a weak attractor network, so that the target-selective population, while in the high memory state, does not strongly suppress distractor cells. When the distractor population has higher activity than target cells in PPC, the locus of attention is transiently shifted to the distractor location (Bisley and Goldberg, 2003, 2006). Representation of the distractor in PPC is thought to be functionally desirable, as it allows the PPC to flexibly function as a saliency map (Bisley and Goldberg, 2010).

After the distractor stimulus is withdrawn, an interesting dynamic occurs in PPC: feedback from the PFC switches the PPC back to encoding the target. This switch back to encoding the target in PPC is a feature of distributed processing in the fronto-parietal circuit. In an isolated local attractor network, a strong distractor response would switch the state of the network from encoding the target to encoding the distractor even after withdrawal of the distractor stimulus (Fig. 2A) (Brunel and Wang, 2001; Compte et al., 2000). In the distributed circuit model, this switch back to encoding the target in PPC is accomplished by feedback projections from PFC. The target cells in PFC send excitation to the target population in PPC and inhibition to the distractor population in PPC. Because PPC is a weak attractor network, this combination of same-selectivity excitation and cross-selectivity inhibition from PFC can effectively switch PPC back to the target memory state. Single-neuron recordings in LIP have shown that PPC networks can switch back to encoding the target in memory after transiently but strongly encoding the distractor (Bisley and Goldberg, 2003; Falkner et al., 2010; Suzuki and Gottlieb, 2013). Furthermore, and consistent with the model, there is experimental evidence that long-range projections between PFC and PPC produce both enhancement and suppression of activity of the recipient cells (Chafee and Goldman-Rakic, 2000).

The PFC exhibits markedly different activity than the PPC in response to distractors. The PFC is activated indirectly, via the PPC’s response to the stimulus. As with the PPC, information about the target is maintained in the PFC by persistent activity in the target-selective population. Subsequent distractor presentation causes a transient suppression of delay activity the PFC. The transient suppression of the target population in PFC is primarily attributable to feedforward different-selectivity inhibition from PPC, rather than local lateral inhibition from the distractor population in PFC. This is consistent with the finding of Suzuki and Gottlieb (2013) that target-selective neurons in the PFC can show suppression by the distractor stimulus without distractor-selective PFC neurons being strongly activated. In contrast to the PPC, the distractor is not represented strongly in the PFC (see also Fig. 4A, bottom). This is because the PFC module is modeled as a strong attractor network, with increased local structure relative to the PPC. This local structure, combining self-excitation and cross-inhibition, allows the target population in the memory state to maintain higher activity than the distractor population throughout the delay. This is the mechanism for the robustness of WM against distraction in the circuit.

We investigated the behavior of the PPC-PFC circuit during error trials. An error trial occurs when the PFC module fails to represent the target at the time of decision or readout, following the delay period (at *t* > 3000 ms). During an error trial, the distractor population fires at a higher rates than the target population (Fig. 5 *B*). This also corresponds to the locus of attention being shifted to the distractor location (Bisley and Goldberg, 2003, 2006). Fig. 5*C* shows that the proportion of errors, and thus distractibility, decreases with greater time separation between target and distractor onset, a behavioral feature that is in line with experimental findings (Suzuki and Gottlieb, 2013). As with the timing dependence of the distractor amplitude in the PPC (Fig. 4*A*), this time course of distractibility is due to the transient dynamics of synaptic gating variables as they evolve toward a steady-state attractor state encoding the target.

### Prefrontal inactivation impairs robustness against distractors

In our model, we simulated inactivation or lesion of the PFC by removing the PFC!PPC feedback inputs to the PPC, and characterized its effects on neural activity and robustness of WM against distractors (Fig. 5 *C,D*). As shown in Fig. 5*D*, PFC inactivation renders the system vulnerable to distractors. Since the PPC exhibits attractor dynamics as an independent local network, the PPC can still encode the target into the delay period through persistent activity, providing some capacity for WM with only PPC engagement. However, without feedback from PFC, the distractor stimulus switches the network to encoding the distractor, which it continues to encode through the subsequent delay. This is in line with experiments finding that PFC inactivation induces error responses to the distractor location (Suzuki and Gottlieb, 2013). This demonstrates the key role of PFC!PPC feedback projections, in the intact circuit, in switching the PPC network back to encoding the target following distractor withdrawal (Fig. 5A). The resulting behavioral vulnerability to distractors is exemplified by an increase in the error rate with respect to control (Fig. 5C). That PFC is not essential for simple WM maintenance — but plays a key role in robustness of WM and filtering of distractors — is in line with conclusions from a range of experimental findings, both in monkeys (Suzuki and Gottlieb, 2013) and in humans (Sakai et al., 2002; Feredoes et al., 2011).

The model findings suggest two modes of operation during WM. When PFC is engaged, the network can operate in a ‘remember first’ regime, storing the initial stimulus and filtering subsequent stimuli. When PFC is not engaged, PPC on its own operates in a ‘remember last’ regime, storing the location of the most recent stimulus, which may also be functionally desirable for a saliency map (Bisley and Goldberg, 2010). As previously mentioned, the amplitude of the transient response to the distractor in PPC is lower compared to that of the target. When the PFC is engaged, the peak distractor response is suppressed (Fig. 5 *A*, top) as compared to when the PFC is inactivated (Fig. 5 *D*). Along these lines, we suggest that the effective strength of lateral inhibition in PPC is not a purely local property and can be flexibly controlled by top-down, i.e., feedback, prefrontal engagement (Falkner et al., 2010, 2013). Our WM modeling results relate to the tradeoff introduced previously (Fig. 2) in that it is possible to have two modules with different recurrent strengths with the capability of filtering out distractors as a unified system.

### Evidence accumulation during perceptual decision making

Having characterized differential roles in WM for the two modules of our distributed circuit, we now examine how it performs in perceptual DM tasks. As shown above in Fig. 2, the recurrent structure *J*_*S*_ of a local network shapes how it accumulates perceptual evidence over time to guide a decision, which suggests the PPC and PFC modules will show differential responses to accumulated evidence during perceptual DM. To probe this issue, we adapted a task paradigm used to study perceptual integration in the mouse (Hanks et al., 2015; Brunton et al., 2013). Hanks et al. (2015) found that in the mouse cortex, frontal and parietal areas differ in their representations of accumulated evidence: parietal neurons encoded the accumulator value in a graded fashion, whereas prefrontal neurons encoded the accumulator value more categorically.

In the two-alternative forced choice paradigm of Hanks et al. (2015), the subject receives auditory input consisting of a sequence of Poisson-distributed clicks each emitted from the left and from the right side, and the subject is rewarded for reporting which side (left vs. right) had the higher frequency signal (Brunton et al., 2013; Hanks et al., 2015). We model clicks as current pulses whose onset is represented by a set of Poisson-distributed times for each trial, where each trial is characterized by a rate pair sorted according to difficulty. For instance, a 18 clicks/s (left): 16 clicks/s (right) trial is “hard” whereas a 30:4 trial is “easy”. The click-triggered current pulses define the inputs to a theoretical accumulator as well as to the circuit model via the first module, the PPC (see Materials and Methods for details).

Fig. 6*A*, top, shows the accumulator value as a function of time, for example trials of varying difficulty. Positive accumulator values correspond to the preferred tuning direction of a neuron, and negative values correspond to the non-preferred direction. The difficulty of the trials is reflected on the slope of the accumulator vs. time plot, where a higher or lower slope in absolute value corresponds to an easy or hard trial, respectively. The trial-averaged firing rates of the PPC and PFC as a function of time and difficulty are shown in Figures 6*A*, middle, and bottom, respectively. The difficulty of the trials is also reflected in the instantaneous slopes of the firing rate, but due to the attractor dynamics and the coupling between the modules, the firing rates in the PPC and PFC are not as linear as a perfect accumulator.

To examine how the theoretical accumulator value is represented in the firing activity of a neural population, we obtained an explicit relationship between accumulator value and firing rate, following the approach of Hanks et al. (2015) (see Materials and Methods). The relationship between firing rate and accumulator value as a function of time is relatively stable for both the PPC and PFC (Fig. 6*C*, top and bottom), similar to empirical findings (Hanks et al., 2015). Importantly, the spacing between different accumulator values is more uniform in the PPC, reflecting a more quasi-linear encoding of the accumulator value as compared to the PFC. We obtained the global relationship between firing rate in the PPC and PFC and accumulator by averaging the plots in Fig. 6*C* with respect to time and scaling the ranges from 0 to 1 (Fig. 6 *B*). We found that, although the accumulator value is encoded in the firing rate for both the PPC and PFC, the encoding is more categorical in PFC versus PPC, as found empirically by Hanks et al. (2015). Indeed, trajectories in the phase plane for a strong attractor (PFC) are shorter than for a weak attractor (PPC). This dynamical difference manifests itself as a steeper slope for the PFC’s firing rate as a function of both time and the hypothetical accumulator which scales linearly with time. These results suggest that the differences in recurrent structure between parietal and prefrontal circuits may contribute to the differences in accumulator encoding between these cortical regions.

### Perceptual decision making across functional cell types

In the primate, the PPC and PFC are key cortical areas engaged in perceptual DM tasks such as visual search and target selection (Schall and Thompson, 1999; Thomas and Pare,´ 2007; Purcell et al., 2010). Within fronto-parietal circuits, saccadic target selection involves at least two stages of processing: selection or discrimination of the relevant target among potential distractors, and preparation of an action or response following that selection (Schall and Thompson, 1999; Woodman et al., 2008). Single-neuron recordings have found that signals related to these two stages are represented heterogeneously across different functional cell types which are distributed across PFC and PPC circuits. Within the frontal eye field (FEF) and other areas, two broad types of neurons — visual cells and movement cells — show distinct dynamics during visual target selection tasks.

The dynamics of visual and movement cells appears to reflect the processes perceptual selection and action preparation, respectively (Schall, 2015). Visual cells respond strongly at stimulus onset. After the initial visual transient, they distinguish target from distractor through higher firing rates. Visual selection cells have been characterized in LIP (Thomas and Paré, 2007; Ipata et al., 2006), FEF (Thompson et al., 1996; Sato et al., 2001; Sato and Schall, 2003) and the subcortical superior colliculus (McPeek and Keller, 2002; White and Munoz, 2011). In contrast, movement cells are not activated by stimulus on-set, but their response is tied to saccade onset. Their activity ramps in motor preparation, with an apparent firing-rate threshold that drives a saccade to their associated movement field (Hanes and Schall, 1996; Woodman et al., 2008). Movement cells are also found in FEF Hanes and Schall (1996); Hanes et al. (1998); Woodman et al. (2008) (McPeek and Keller, 2002) and superior colliculus, but appear to be much more sparse in LIP (Ferraina et al., 2002).

We sought to test whether our distributed circuit model could capture these differences between visual and movement functional cells in fronto-parietal circuits. In the context of saccadic target selection, we associate Module 1 with Selection (visual) cells and Module 2 with Action (movement) cells, and these functional cell types may be differentially distributed across PPC and PFC. Sensory input enters into Selection cells (Module 1), as proposed for a sensorimotor cascade (Schall, 2013; Purcell et al., 2010). Choice and reaction time is set by a firing-rate threshold on Action cells (Module 2) (Hanes and Schall, 1996; Woodman et al., 2008). The goal of the visual target selection task is to make a saccade towards a target in the presence of distractors. The difficulty of the task is dictated by the contrast, which reflects the salience of the target with respect to that of distractors (Thomas and Paré, 2007; Sato et al., 2001). High or low contrast corresponds to low or high target-distractor similarity, respectively.

In the model, stimulus input activates Selection cells (Fig. 7*A*). When the stimulus appears, there is a pronounced visual transient in the firing rate of both target and distractor populations in the Selection module. The target-selective population receives more input than the distractor population (the amount depending on the contrast, see Eq. 14), and the activity of the two populations diverge due to competitive dynamics (Wong and Wang, 2006), resulting in a discrimination of the target from the distractor. Due to the feedforward projection from the Selection module to the Action module, Action cells receive stimulus signals indirectly. In a correct trial, the target population reaches the response threshold and a reaction time is recorded. In a high-contrast condition, the reaction time is lower because there is a larger difference in input current to the target and distractor populations of the Selection module (Wong and Wang, 2006). Thus, the contrast-dependent differences in input current to the populations in the Selection module along with the amplification of those differences due to the recurrent dynamics of both modules eventually leads to a categorical choice in the Action cells.

The high-activity population, corresponding to the choice, is consistent in both Selection and Action modules, but there are two important differences in the dynamics between the modules. First, the competitive dynamics in the Action module, as compared to the Selection module, result in a steeper firing-rate ramping as a function of time (Fig. 7*A*). This is because the Action module has a higher recurrent structure **J*_S_*, and stronger attractor dynamics, than the Selection module. Weaker structure in the Selection module enables better integration of perceptual evidence for target selection, as shown in Fig. 2.

Second, the pronounced transient activation of Selection cells, due to the appearance of the target, is not represented in the Action module (Fig. 7*A*). This is due to the pathway-specific excitation-inhibition (E/I) balance in the feedforward Selection→Action projection (Eq. 8). Therefore net inputs to the Action module reflect the difference of activity between the populations in the Selection module. Thus, an Action-cell response above baseline will only be observed when the activities in the Selection module have diverged, in line with theories and evidence of “discrete flow” between selection-and action-related stages in perceptual DM (Woodman et al., 2008). These results suggest that the functional distinction between Selection and Action cells arises from a difference in structure in the respective modules and the existence of pathway-specific E/I balance onto Action cells.

Single-neuron recordings during visual search have characterized how dynamics of functional cell types relate to reaction times, through the target-distractor discrimination time in visual cells, and onset time in movement cells (see Materials and Methods). For visual cells, the target-distractor discrimination time correlates with reaction time, across and within search difficulty conditions (Sato et al., 2001; Thomas and Paré, 2007; Ipata et al., 2006; McPeek and Keller, 2002; White and Munoz, 2011). For movement cells, the onset time correlates with reaction time (Woodman et al., 2008; McPeek and Keller, 2002; White and Munoz, 2011). We computed analogous measures for Selection and Action cells in our model. In line with experimental findings, we found that reaction time correlates with discrimination time in Selection cells and with onset time in Action cells, across different contrast conditions as well as across reaction-time variability within a contrast condition (Fig. 7B). This implies that although selection and action are distinct stages in perceptual DM, they are consistent and reflect the feedforward nature of the two-module circuit architecture.

### Pathway-specific E/I balance and speed-accuracy tradeoff

We have shown that Action cells are activated only after the Selection cells have diverged to select an option, because the projection from Selection to Action cells exhibits pathway-specific E/I balance. To further characterize the role of pathway-specific E/I balance in the feedforward projection from Selection to Action cells, we parametrically reduced the strength of feedforward inhibition while holding constant the strength of feed-forward excitation (Eq. 15). For each level of inhibition, we computed a chronometric and a psychometric plot: reaction time as a function of contrast and accuracy as a function of contrast (Fig. 8*A*). As feedforward inhibition decreases, reaction times decrease, but so does accuracy. Both of these effects are more pronounced at lower contrast values. Therefore, perturbing pathway-specific E/I balance implements a speed-accuracy tradeoff during target selection.

We then examined how errors arise in the model under control balanced and reduced-inhibition conditions. Fig. 8B shows representative single trials from the two conditions. In the control balanced condition, all errors arise due to mis-selection of the distractor instead of the target in Selection cells (Fig. 8*B*, left), which is in line with single-neuron recordings finding mis-selection in visual cells during search tasks (Thompson et al., 2005; Trageser et al., 2008; Shen and Pare,´ 2007). In contrast, under reduced inhibi-tion a new type of error trial can occur (Fig. 8*B*, right). Action cells prematurely select a response before target selection has completed within the Selection cells, causing errors because the Action module makes a decision for target or distractor in a quasi-random manner. Under reduced inhibition, when the target and distractor Selection cells are activated, but not yet diverged, the Action module receives a non-specific net-excitatory input. This net-excitatory input can induce quasi-random winner-take-all DM (Wong and Wang, 2006), giving rise to an imbalance-dependent type of error. Our findings suggest that in a healthy physiological state, the projection from Module-1 cells to Module-2 cells should be in a state near E/I balance, because this configuration produces errors consistent with electrophysiological recordings (Thompson et al., 2005; Trageser et al., 2008; Shen and Paré, 2007).

## Discussion

In this study, we propose a parsimonious circuit model for distributed computation subserving WM and DM, two core cognitive functions that recruit overlapping fronto-parietal circuits. We highlight the tradeoff that exists when optimizing recurrent structure for robustness against distractors in WM versus slow integration of evidence in DM; the model developed here ameliorates this tradeoff by extending the local circuit to two modules. We found that across both WM and DM paradigms, the circuit model captures a wide range of salient, empirically observed features of neural activity in fronto-parietal circuits. We summarize the model’s findings with respect to the circuit architecture and its relationship to WM and DM computations.

First, Module 1 (PPC or Selection cells), which receives sensory input (Felleman and Van Essen, 1991; Buschman and Miller, 2007; Ibos et al., 2013; Siegel et al., 2015), is a weak attractor network. This property is beneficial so that PPC can transiently encode distractors and function as a saliency map, and so that its memory state can be effectively controlled by weak feedback projections from PFC (e.g., to switch the state back to encoding the target following distractor presentation). In the context of perceptual DM, Module 1 (Selection cells) should be a weak attractor network to support integration of perceptual evidence with a long timescale to improve accuracy.

Second, Module 2 (PFC or Action cells) is a strong attractor network. In the context of WM, this property is functionally beneficial because PFC can thereby provide robustness against distractors, filtering out the effects of strong distractor responses in PPC. In the context of DM, it is functionally beneficial for Action cells to be a strong attractor network because this enables them to ramp quickly to threshold to drive choice, following the upstream target selection. This difference in recurrent structure also predicts a difference in the representation of accumulated value, which is more graded in Module 1 and more categorical in Module 2.

Third, the modules are interconnected via reciprocal projections that are structured: net-excitatory between same-selectivity populations and net-inhibitory between different-selectivity populations. The feedforward Module 1→Module 2 projection should be structured to propagate signals for both WM and DM. The feedback Module 2→Module 1 projection is especially important in the context of WM. This projection being structured allows PFC to switch PPC back to encoding the target following distractor presentation.

### Differential roles for PPC and PFC in working memory

Although PPC and PFC are both involved in active WM maintenance, converging evidence from a range of methodologies suggests differential roles, with PPC associated with attentional saliency and selection (Wardak et al., 2012, 2002), and PFC associated with robustness of WM and filtering of distractors (Suzuki and Gottlieb, 2013; Sakai et al., 2002; Feredoes et al., 2011). Our distributed WM circuit model captures multiple key features from these studies and suggests that the different roles of PPC and PFC may in part be due to distinct dynamical behaviors arising from their recurrent microcircuitry (Chaudhuri et al., 2015), as suggested by electrophysiological recordings (Murray et al., 2014b; Katsuki et al., 2014).

The model also proposes a key role for feedback from PFC to PPC during distractor processing, namely to strengthen target representations via feedback excitation and filter distractor representations via feedback inhibition (Figs. 4,5). In line with this proposal, Feredoes et al. (2011) performed combined stimulation-imaging experiments in humans and found a key role for feedback from PFC to posterior areas during distractor filtering to enhance target representations (see also Edin et al. (2009) for PFC-mediated enhancement of WM capacity in PPC). Recording in LIP during visuospatial WM, Falkner et al. (2010, 2013) found that surround suppression of distractors, and target representation, are strengthened by top-down cognitive modulation (e.g., by motivation). Our model predicts that PFC inactivation should disrupt this modulation of surround suppression in PPC. Furthermore, if PFC is inactivated or the PFC-PFC feedback projection is very weak, the system operates in a ‘remember-last’ regime (Fig. 5*D*) whereas if PFC is engaged the system operates in a ‘remember-first’ regime (Fig. 5*A*).

### Evidence accumulation in the fronto-parietal network

Gradual accumulation of perceptual evidence is reflected in DM-related neuronal activity in fronto-parietal circuits (Gold and Shadlen, 2007; Schall, 2013; Brody and Hanks, 2016). Drift-diffusion models (Purcell et al., 2010; Bogacz et al., 2006; Usher and McClelland, 2001) as well as recurrent circuit models (Wang, 2002; Wong and Wang, 2006) can account for the accumulation process and subsequent ramping behavior in the neural dynamics, including more discrete “jumping” modes (Miller and Katz, 2010; Wang, 2012; Lo et al., 2015; Latimer et al., 2015). Our model predicts that, due to the inter-areal differences in local recurrent structure, representations of accumulated evidence are more graded in PPC and more categorical in PFC, as supported by single-neuron recordings (Hanks et al., 2015). Recent inactivation studies found that even though PPC shows DM-related signals, it does not play a causal role in perceptual DM (Erlich et al., 2015; Katz et al., 2016). It is unclear whether these results can be accounted for via compensatory mechanisms and/or distributed processing within a broader region in PPC that includes LIP. In rats, inactivation of PPC disrupts accumulation of visual but not auditory evidence (Raposo et al., 2014). In monkeys, inactivation of LIP disrupts saccadic selection of visuospatial targets during visual search tasks (Wardak et al., 2002, 2004), which are more similar to the DM paradigm we have modeled.

### Dynamics and localization of visual and movement cells

In the context of perceptual DM, the Selection (Module 1) and Action (Module 2) cells in our model capture key dynamical features of visual and movement cells, respectively, which have been characterized in single-neuron recordings from LIP, FEF, and SC (Schall, 2015; Sato et al., 2001; Thompson et al., 1996; Thomas and Paré, 2007; Ipata et al., 2006; Hanes et al., 1998; Woodman et al., 2008; McPeek and Keller, 2002). Visual and movement cells in cortex may have distinct biophysical properties (Cohen et al., 2009b), laminar distributions (Pouget et al., 2009), and long-range projections (Gregoriou et al., 2012). Our results suggest that both modules could represent neural populations that are distributed across multiple areas (e.g., visual cells in LIP/FEF/SC, movement cells in FEF/SC).

An important feature of our distributed circuit model is the pathway-specific E/I balance in the feedforward projection from Selection to Action cells. This E/I balance regulates a cascade of activations across functional cell types and can be characterized as “discrete flow,” since response preparation in Action cells only begins after target selection is completed in Selection cells (Woodman et al., 2008). Pathway-specific balance thereby provides a mechanism for discrete flow without requiring direct gating of neuronal responses (Purcell et al., 2010; Wang et al., 2004; Yang et al., 2016). The sensitivity of Action cells to differences in upstream Selection cells is related to a geometrical characterization of neural dynamics, whereby some directions in neural state space elicit responses while those in the null-space do not (Vogels and Abbott, 2009; Li et al., 2016; Kaufman et al., 2014).

We disrupted pathway-specific E/I balance by systematically decreasing the strength of feedforward inhibition onto the Action cells. A strongly imbalanced circuit produced a distinct type of error: Action cells started ramping before the populations of Selection cells had diverged in activity, i.e., before selection was accomplished. The imbalanced condition may capture dynamics recorded from visual and movement cells in FEF under speed-demanding compelled response paradigms (Stanford et al., 2010). The modulation of feedforward inhibition in our model resulted in a smooth tradeoff between speed and accuracy, a plausible mechanism among others (Heitz and Schall, 2012; Bogacz et al., 2010; Standage et al., 2014; Hanks et al., 2015; Stanford et al., 2010).

### Limitations and future directions

Future studies can build upon and extend the present model to address a number of important questions. One direction is to extend the two-population discrete network studied here to a quasi-continuous network in which neurons exhibit smoothly varying tuning of a parametric stimulus variable (Compte et al., 2000; Furman and Wang, 2008) to explore effects that depend on the similarity and distance between distractors and targets held in WM (Suzuki and Gottlieb, 2013; Murray et al., 2014a). Generalization to more than two populations would enable modeling of set-size effects in visual search tasks, whereby the number of stimuli affects behavior and visual- and movement-cell activity (Cohen et al., 2009a; Balan et al., 2008; Woodman et al., 2008). Extension to a spiking circuit model would enable modeling the neural signatures of synchronization between PPC and PFC during cognitive processing (Ardid et al., 2010; Pesaran et al., 2008; Salazar et al., 2012; Dotson et al., 2014).

This study suggests important design principles for constructing multi-regional neural circuit models of distributed cognitive function, such as the interplay between long-range and local connectivity in recurrent dynamics and computation, the roles of specialized microcircuit properties across the cortical hierarchy, and the implications of balanced excitation and inhibition in long-range interactions. The parsimonious model studied here may therefore instantiate features of a canonical cognitive circuit useful for studying distributed computation in the brain.

## References

Abbott LF, Chance FS (2005) Drivers and modulators from push-pull and balanced synaptic input. Prog Brain Res 149:147–55.

Amit DJ, Brunel N (1997) Model of global spontaneous activity and local structured activity during delay periods in the cerebral cortex. Cereb Cortex 7:237–252.

Ardid S, Wang XJ, Gomez-Cabrero D, Compte A (2010) Reconciling coherent oscillation with modulation of irregular spiking activity in selective attention: gamma-range synchronization between sensory and executive cortical areas. J Neurosci 30:2856–70.

Balan PF, Oristaglio J, Schneider DM, Gottlieb J (2008) Neuronal correlates of the set-size effect in monkey lateral intraparietal area. PLoS Biol 6:e158.

Bisley JW, Goldberg ME (2003) Neuronal activity in the lateral intraparietal area and spatial attention. Science 299:81–6.

Bisley JW, Goldberg ME (2006) Neural correlates of attention and distractibility in the lateral intraparietal area. J Neurophysiol 95:1696–717.

Bisley JW, Goldberg ME (2010) Attention, intention, and priority in the parietal lobe. Annu Rev Neurosci 33:1–21.

Bogacz R, Brown E, Moehlis J, Holmes P, Cohen JD (2006) The physics of optimal decision making: a formal analysis of models of performance in two-alternative forced-choice tasks. Psychol Rev 113:700–65.

Bogacz R, Wagenmakers EJ, Forstmann BU, Nieuwenhuis S (2010) The neural basis of the speed-accuracy tradeoff. Trends Neurosci 33:10–6.

Brody CD, Hanks TD (2016) Neural underpinnings of the evidence accumulator. Curr Opin Neurobiol 37:149–57.

Brunel N, Wang XJ (2001) Effects of neuromodulation in a cortical network model of object working memory dominated by recurrent inhibition. Journal of computational neuroscience 11:63–85.

Brunton BW, Botvinick MM, Brody CD (2013) Rats and humans can optimally accumulate evidence for decision-making. Science 340:95–8.

Buschman TJ, Miller EK (2007) Top-down versus bottom-up control of attention in the prefrontal and posterior parietal cortices. Science 315:1860–2.

Chafee MV, Goldman-Rakic PS (1998) Matching patterns of activity in primate pre-frontal area 8a and parietal area 7ip neurons during a spatial working memory task. J Neurophysiol 79:2919–40.

Chafee MV, Goldman-Rakic PS (2000) Inactivation of parietal and prefrontal cortex reveals interdependence of neural activity during memory-guided saccades. J Neurophysiol 83:1550–66.

Chaudhuri R, Knoblauch K, Gariel MA, Kennedy H, Wang XJ (2015) A large-scale circuit mechanism for hierarchical dynamical processing in the primate cortex. Neuron 88:419–31.

Christophel TB, Klink PC, Spitzer B, Roelfsema PR, Haynes JD (2017) The distributed nature of working memory. Trends in Cognitive Sciences 21:111–124.

Clewley R (2012) Hybrid models and biological model reduction with PyDSTool. PLoS Comput Biol 8:e1002628.

Cohen JY, Heitz RP, Woodman GF, Schall JD (2009a) Neural basis of the set-size effect in frontal eye field: timing of attention during visual search. J Neurophysiol 101:1699–704.

Cohen JY, Pouget P, Heitz RP, Woodman GF, Schall JD (2009b) Biophysical support for functionally distinct cell types in the frontal eye field. J Neurophysiol 101:912–6.

Compte A, Brunel N, Goldman-Rakic PS, Wang XJ (2000) Synaptic mechanisms and network dynamics underlying spatial working memory in a cortical network model. Cereb Cortex 10:910–23.

Constantinidis C, Procyk E (2004) The primate working memory networks. Cogn Affect Behav Neurosci 4:444–465.

Constantinidis C, Wang XJ (2004) A neural circuit basis for spatial working memory. Neuroscientist 10:553–65.

Domenech P, Redoute´ J, Koechlin E, Dreher JC (2017) The neuro-computational architecture of value-based selection in the human brain. Cerebral Cortex.

Dotson NM, Salazar RF, Gray CM (2014) Frontoparietal correlation dynamics reveal interplay between integration and segregation during visual working memory. J Neurosci 34:13600–13.

Duncan J (2010) The multiple-demand (md) system of the primate brain: mental programs for intelligent behaviour. Trends Cogn Sci 14:172–9.

Edin F, Klingberg T, Johansson P, McNab F, Tegner´ J, Compte A (2009) Mechanism for top-down control of working memory capacity. Proceedings of the National Academy of Sciences 106:6802–6807.

Erlich JC, Brunton BW, Duan CA, Hanks TD, Brody CD (2015) Distinct effects of prefrontal and parietal cortex inactivations on an accumulation of evidence task in the rat. Elife 4.

Everling S, Tinsley CJ, Gaffan D, Duncan J (2002) Filtering of neural signals by focused attention in the monkey prefrontal cortex. Nat Neurosci 5:671–6.

Falkner AL, Goldberg ME, Krishna BS (2013) Spatial representation and cognitive modulation of response variability in the lateral intraparietal area priority map. J Neurosci 33:16117–30.

Falkner AL, Krishna BS, Goldberg ME (2010) Surround suppression sharpens the priority map in the lateral intraparietal area. J Neurosci 30:12787–97.

Felleman DJ, Van Essen DC (1991) Distributed hierarchical processing in the primate cerebral cortex. Cereb Cortex 1:1–47.

Feredoes E, Heinen K, Weiskopf N, Ruff C, Driver J (2011) Causal evidence for frontal involvement in memory target maintenance by posterior brain areas during distracter interference of visual working memory. Proc Natl Acad Sci U S A 108:17510–5.

Ferraina S, Pare´ M, Wurtz RH (2002) Comparison of cortico-cortical and cortico-collicular signals for the generation of saccadic eye movements. J Neurophysiol 87:845–58.

Funahashi S, Bruce CJ, Goldman-Rakic PS (1989) Mnemonic coding of visual space in the monkey’s dorsolateral prefrontal cortex. J Neurophysiol 61:331–49.

Furman M, Wang XJ (2008) Similarity effect and optimal control of multiple-choice decision making. Neuron 60:1153–68.

Gold JI, Shadlen MN (2007) The neural basis of decision making. Annu Rev Neurosci 30:535–74.

Goldman-Rakic PS (1995) Cellular basis of working memory. Neuron 14:477–85.

Gregoriou GG, Gotts SJ, Desimone R (2012) Cell-type-specific synchronization of neural activity in fef with v4 during attention. Neuron 73:581–94.

Hanes DP, Patterson n WF, Schall JD (1998) Role of frontal eye fields in countermanding saccades: visual, movement, and fixation activity. J Neurophysiol 79:817–34.

Hanes DP, Schall JD (1996) Neural control of voluntary movement initiation. Science 274:427–30.

Hanks TD, Kopec CD, Brunton BW, Duan CA, Erlich JC, Brody CD (2015) Distinct relationships of parietal and prefrontal cortices to evidence accumulation. Nature 520:220–3.

Heitz RP, Schall JD (2012) Neural mechanisms of speed-accuracy tradeoff. Neuron 76:616–28.

Ibos G, Duhamel JR, Ben Hamed S (2013) A functional hierarchy within the parietofrontal network in stimulus selection and attention control. Journal of Neuroscience 33:8359–8369.

Ipata AE, Gee AL, Goldberg ME, Bisley JW (2006) Activity in the lateral intraparietal area predicts the goal and latency of saccades in a free-viewing visual search task. J Neurosci 26:3656–61.

Katsuki F, Constantinidis C (2012a) Early involvement of prefrontal cortex in visual bottom-up attention. Nat Neurosci 15:1160–6.

Katsuki F, Constantinidis C (2012b) Unique and shared roles of the posterior parietal and dorsolateral prefrontal cortex in cognitive functions. Front Integr Neurosci 6:17.

Katsuki F, Qi XL, Meyer T, Kostelic PM, Salinas E, Constantinidis C (2014) Differences in intrinsic functional organization between dorsolateral prefrontal and posterior parietal cortex. Cereb Cortex 24:2334–49.

Katz LN, Yates JL, Pillow JW, Huk AC (2016) Dissociated functional significance of decision-related activity in the primate dorsal stream. Nature 535:285–8.

Kaufman MT, Churchland MM, Ryu SI, Shenoy KV (2014) Cortical activity in the null space: permitting preparation without movement. Nat Neurosci 17:440–8.

Latimer KW, Yates JL, Meister MLR, Huk AC, Pillow JW (2015) Single-trial spike trains in parietal cortex reveal discrete steps during decision-making. Science 349:184–7.

Lawrence BM, White r RL, Snyder LH (2005) Delay-period activity in visual, visuomovement, and movement neurons in the frontal eye field. J Neurophysiol 94:1498–508.

Li N, Daie K, Svoboda K, Druckmann S (2016) Robust neuronal dynamics in premotor cortex during motor planning. Nature 532:459–64.

Lo CC, Wang CT, Wang XJ (2015) Speed-accuracy tradeoff by a control signal with balanced excitation and inhibition. Journal of Neurophysiology 114:650–661.

Machens CK, Romo R, Brody CD (2005) Flexible control of mutual inhibition: a neural model of two-interval discrimination. Science 307:1121–4.

McPeek RM, Keller EL (2002) Saccade target selection in the superior colliculus during a visual search task. J Neurophysiol 88:2019–34.

Meister MLR, Hennig JA, Huk AC (2013) Signal multiplexing and single-neuron computations in lateral intraparietal area during decision-making. J Neurosci 33:2254–67.

Miller EK, Erickson CA, Desimone R (1996) Neural mechanisms of visual working memory in prefrontal cortex of the macaque. J Neurosci 16:5154–5167.

Miller EK, Li L, Desimone R (1993) Activity of neurons in anterior inferior temporal cortex during a short-term memory task. J Neurosci 13:1460–78.

Miller P, Katz DB (2010) Stochastic transitions between neural states in taste processing and decision-making. J Neurosci 30:2559–70.

Mitchell DJ, Bell AH, Buckley MJ, Mitchell AS, Sallet J, Duncan J (2016) A putative multiple-demand system in the macaque brain. J Neurosci 36:8574–85.

Murray JD, Anticevic A, Gancsos M, Ichinose M, Corlett PR, Krystal JH, Wang XJ (2014a) Linking microcircuit dysfunction to cognitive impairment: effects of disinhibition associated with schizophrenia in a cortical working memory model. Cereb Cortex 24:859–72.

Murray JD, Bernacchia A, Freedman DJ, Romo R, Wallis JD, Cai X, Padoa-Schioppa C, Pasternak T, Seo H, Lee D, Wang XJ (2014b) A hierarchy of intrinsic timescales across primate cortex. Nat Neurosci 17:1661–3.

Pesaran B, Nelson MJ, Andersen RA (2008) Free choice activates a decision circuit between frontal and parietal cortex. Nature 453:406–9.

Pouget P, Stepniewska I, Crowder EA, Leslie MW, Emeric EE, Nelson MJ, Schall JD (2009) Visual and motor connectivity and the distribution of calcium-binding proteins in macaque frontal eye field: implications for saccade target selection. Front Neuroanat 3:2.

Powell KD, Goldberg ME (2000) Response of neurons in the lateral intraparietal area to a distractor flashed during the delay period of a memory-guided saccade. J Neurophysiol 84:301–10.

Purcell BA, Heitz RP, Cohen JY, Schall JD, Logan GD, Palmeri TJ (2010) Neurally constrained modeling of perceptual decision making. Psychol Rev 117:1113–43.

Qi XL, Katsuki F, Meyer T, Rawley JB, Zhou X, Douglas KL, Constantinidis C (2010) Comparison of neural activity related to working memory in primate dorsolateral pre-frontal and posterior parietal cortex. Front Syst Neurosci 4:12.

Raposo D, Kaufman MT, Churchland AK (2014) A category-free neural population supports evolving demands during decision-making. Nat Neurosci 17:1784–92.

Ratcliff R (1978) A theory of memory retrieval. Psychological Review 85:59–108.

Sakai K, Rowe JB, Passingham RE (2002) Active maintenance in prefrontal area 46 creates distractor-resistant memory. Nat Neurosci 5:479–84.

Salazar RF, Dotson NM, Bressler SL, Gray CM (2012) Content-specific fronto-parietal synchronization during visual working memory. Science 338:1097–100.

Sato T, Murthy A, Thompson KG, Schall JD (2001) Search efficiency but not response interference affects visual selection in frontal eye field. Neuron 30:583–91.

Sato TR, Schall JD (2003) Effects of stimulus-response compatibility on neural selection in frontal eye field. Neuron 38:637–48.

Schall JD (2001) Neural basis of deciding, choosing and acting. Nat Rev Neurosci 2:33–42.

Schall JD, Thompson KG (1999) Neural selection and control of visually guided eye movements. Annu Rev Neurosci 22:241–59.

Schall JD (2013) Macrocircuits: decision networks. Curr Opin Neurobiol 23:269–74.

Schall JD (2015) Visuomotor functions in the frontal lobe. Annual Review of VisionScience 1:469–498.

Shen K, Pare´ M (2007) Neuronal activity in superior colliculus signals both stimulus identity and saccade goals during visual conjunction search. J Vis 7:15.1–13.

Siegel M, Buschman TJ, Miller EK (2015) Cortical information flow during flexible sensorimotor decisions. Science 348:1352–5.

Standage D, Blohm G, Dorris MC (2014) On the neural implementation of the speed-accuracy trade-off. Front Neurosci 8:236.

Standage D, Pare´ M (2011) Persistent storage capability impairs decision making in a biophysical network model. Neural Netw 24:1062–73.

Stanford TR, Shankar S, Massoglia DP, Costello MG, Salinas E (2010) Perceptual decision making in less than 30 milliseconds. Nat Neurosci 13:379–85.

Suzuki M, Gottlieb J (2013) Distinct neural mechanisms of distractor suppression in the frontal and parietal lobe. Nat Neurosci 16:98–104.

Thomas NWD, Pare´ M (2007) Temporal processing of saccade targets in parietal cortex area LIP during visual search. J Neurophysiol 97:942–7.

Thompson KG, Hanes DP, Bichot NP, Schall JD (1996) Perceptual and motor processing stages identified in the activity of macaque frontal eye field neurons during visual search. J Neurophysiol 76:4040–55.

Thompson KG, Bichot NP, Sato TR (2005) Frontal eye field activity before visual search errors reveals the integration of bottom-up and top-down salience. J Neurophysiol 93:337–51.

Trageser JC, Monosov IE, Zhou Y, Thompson KG (2008) A perceptual representation in the frontal eye field during covert visual search that is more reliable than the behavioral report. Eur J Neurosci 28:2542–9.

Usher M, McClelland JL (2001) The time course of perceptual choice: the leaky, competing accumulator model. Psychol Rev 108:550–92.

Vogels TP, Abbott LF (2009) Gating multiple signals through detailed balance of excitation and inhibition in spiking networks. Nat Neurosci 12:483–491.

Wang XJ, Tegner´ J, Constantinidis C, Goldman-Rakic PS (2004) Division of labor among distinct subtypes of inhibitory neurons in a cortical microcircuit of working memory. Proc Natl Acad Sci U S A 101:1368–73.

Wang XJ (1999) Synaptic basis of cortical persistent activity: the importance of NMDA receptors to working memory. J Neurosci 19:9587–9603.

Wang XJ (2001) Synaptic reverberation underlying mnemonic persistent activity. Trends Neurosci 24:455–463.

Wang XJ (2002) Probabilistic decision making by slow reverberation in cortical circuits. Neuron 36:955–968.

Wang XJ (2008) Decision making in recurrent neuronal circuits. Neuron 60:215–34.

Wang XJ (2012) Neural dynamics and circuit mechanisms of decision-making. Current Opinion in Neurobiology 22:1039–1046 Decision making.

Wang XJ (2013) The prefrontal cortex as a quintessential “cognitive-type” neural circuit: working memory and decision making In Stuss DT, Knight RT, editors, Principles of frontal lobe function, chapter 15, pp. 226–248. Oxford University Press, New York, 2nd edition.

Wardak C, Ben Hamed S, Olivier E, Duhamel JR (2012) Differential effects of parietal and frontal inactivations on reaction times distributions in a visual search task. Front Integr Neurosci 6:39.

Wardak C, Olivier E, Duhamel JR (2002) Saccadic target selection deficits after lateral intraparietal area inactivation in monkeys. J Neurosci 22:9877–84.

Wardak C, Olivier E, Duhamel JR (2004) A deficit in covert attention after parietal cortex inactivation in the monkey. Neuron 42:501–8.

White BJ, Munoz DP (2011) Separate visual signals for saccade initiation during target selection in the primate superior colliculus. J Neurosci 31:1570–8.

Wong KF, Huk AC, Shadlen MN, Wang XJ (2007) Neural circuit dynamics underlying accumulation of time-varying evidence during perceptual decision making. Front Comput Neurosci 1:6.

Wong KF, Wang XJ (2006) A recurrent network mechanism of time integration in perceptual decisions. J Neurosci 26:1314–28.

Woodman GF, Kang MS, Thompson K, Schall JD (2008) The effect of visual search efficiency on response preparation: neurophysiological evidence for discrete flow. Psychol Sci 19:128–36.

Yang GR, Murray JD, Wang XJ (2016) A dendritic disinhibitory circuit mechanism for pathway-specific gating. Nat Commun 7:12815.

Zhang W, Falkner AL, Krishna BS, Goldberg ME, Miller KD (2017) Coupling between one-dimensional networks reconciles conflicting dynamics in fLIPg and reveals its recurrent circuitry. Neuron 93:221–234.

